# Niche differences, not fitness differences, explain coexistence across ecological groups

**DOI:** 10.1101/2021.11.15.468654

**Authors:** Lisa Buche, Jurg W. Spaak, Javier Jarillo, Frederik De Laender

**Author notes:** These authors have contributed equally (Lisa Buche, Jurg W. Spaak). **Statement of authorship:** LB and JS designed the collection of data. LB gathered the data, with help from all authors. JS computed the N and F. LB analyzed the data with inputs from JS and FDL. Js performed the clustering. JJ performed the meta-analysis and odd-ratio analysis. LB and JS wrote the first draft of the manuscript, and all authors contributed substantially to revisions. **Data accessibility statement:** Code is available on Github https://github.com/Buchel9844/Meta-analysis_NFD. corresponding authors Lisa Buche.

## Abstract

Understanding the drivers of species coexistence is an important objective in ecology. Yet, the multitude of methods to study coexistence hampers cross-community comparisons. Here, we standardized niche and fitness differences (i.e how species limit themselves compared to others and their competitive ability, respectively) across 1018 species pairs to investigate species coexistence across ecological groups and methodological settings (experimental setup, natural co-occurrence, population model used, and growth method). We find that, first, coexistence is driven by large niche differences, not by small fitness differences. Second, species group into clear clusters of coexisting and non-coexisting species along the niche axis. Finally, these clusters are not driven by ecological or methodological settings. This suggests differences between coexisting and non-coexisting communities transcending those measured in our empirical systems. Overall, our results show that species coexistence is mainly influenced by mechanisms acting on niche differences.

## 1 Introduction

Species interact in a variety of ways. Quantifying the consequences of such interactions for the capacity of species to coexist and form communities is a central objective in ecology (Chesson, 2000b; Barabás, D’Andrea, et al., 2018; Chesson, 2020; Saavedra et al., 2017; J. W. Spaak, Godoy, et al., 2021). One challenge is that, even at a local scale, the diversity of potential mechanisms is overwhelming and difficult to track (Pilosof et al., 2017). At present, we lack a synthetic insight into how the diversity of specific mechanisms, observed across a diversity of ecological groups, maps on to species coexistence. Achieving such a synthesis is challenging because these mechanisms can differ among ecological groups, and is typically analyzed with different methods.

One way to overcome this limitation is by applying the concepts of niche and fitness difference across different ecological groups (Chesson, 2000b; J. W. Spaak and De Laender, 2020; Koffel et al., 2021b; J. W. Spaak, Godoy, et al., 2021). Specifically, niche differences measure how much species limit themselves compared to others, while fitness differences measure how competitive ability (e.g. fecundity, dispersion) varies among species (Chesson, 2000b; Barabás, D’Andrea, et al., 2018). These concepts may then act as a common currency and as such permit synthesis (Grainger, Levine, et al., 2019). While species can physically interact via an unlimited number of mechanisms, information on niche and fitness differences can categorize all these mechanisms into a small number of high level-processes (positive, negative or no frequency dependence, and facilitation), as well as the expected outcomes of these processes (competitive exclusion, coexistence, and priority effects) (J. W. Spaak, Godoy, et al., 2021). Frequency dependence occurs when the population growth rate of a species depends on its relative abundance, with total abundance of all species kept constant (Adler, HilleRisLambers, et al., 2007; Ke and Letten, 2018). When frequency dependence is negative (positive), species grow less well when their relative abundance is high (low). Facilitation occurs when population growth is boosted by the presence of a second species (J. W. Spaak and De Laender, 2020; Bimler et al., 2018). Competitive exclusion (coexistence) occurs when species cannot (can) persist together, irrespective of their initial abundance. Priority effects involve exclusion dependent on initial abundance (Mordecai, 2011; Ke and Letten, 2018).

Niche and fitness differences have been applied in the past to synthesise the causes and consequences of species interactions (e.g.Adler, HilleRisLambers, et al., 2007; Godoy, Kraft, et al., 2014; Godoy and Levine, 2014; Narwani et al., 2013). However, their use comes with an important limitation: there is not a generally accepted method on how to measure niche and fitness differences (Godwin et al., 2020; Song et al., 2019). Rather, there are as many as eleven different methods to assess niche and fitness differences (J. W. Spaak and De Laender, 2020; Koffel et al., 2021a). Many of these methods are tailored to a specific community model and, consequentially, have only been applied to ecological groups which are well described by these models (Bimler et al., 2018; Godoy, 2019). This specificity has hampered cross community comparison. Recently, we developed a method to assess niche and fitness differences (J. W. Spaak and De Laender, 2020; J. W. Spaak, Godoy, et al., 2021) which allows computing niche and fitness differences in a standardized way and/or to convert available data into a common currency. Given this method, it is now possible to use niche and fitness differences as a common currency across multiple ecological groups to ask what permits or hampers species coexistence (Narwani et al., 2013; Germain, Weir, et al., 2016; Grainger, Levine, et al., 2019).

In principle, coexistence can occur when niche differences (promoting coexistence) are large, when fitness differences (hampering coexistence) are small, or when both occur. The importance of the former for coexistence has been demonstrated (mostly in plants) in both semi-natural (Adler, Ellner, et al., 2010; Chu and Adler, 2015; Godoy, Stouffer, et al., 2017; Adler, Smull, et al., 2018; Armitage and Jones, 2019) and experimental communities (Narwani et al., 2013; Mathias and Chesson, 2013; Li Shao-peng et al., 2019). Yet, also small fitness differences have been found to promote coexistence (Chu and Adler, 2015). Thus, it is not clear what explains coexistence: fitness differences, niche differences, or both? A first outstanding question is therefore if, across multiple ecological groups, large niche difference, or small fitness difference, or both, explain coexistence.

Next, we might ask whether coexisting and non-coexisting species differ conceptually in their underlying species interactions. For example, neutrally co-occurring species differ conceptually in their underlying species interactions from stably coexisting species (Hubbell, 2001; Chesson, 2000a), which leads to a gap in their distributions of niche and fitness differences (Song et al., 2019). More generally, if coexisting species differ from non-coexisting species only in the relative strength of the underlying species interactions, then we would expect a gradual change of niche and fitness differences from coexistence to competitive exclusion. Conversely, if coexistence is driven, at least partially, by fundamentally different underlying species interactions, then we except a different distributions of niche or fitness differences for coexisting communities than for non-coexisting communities. A second outstanding question is therefore whether species cluster in the niche differences - fitness differences space.

Here, we performed a metaanalysis of species-interaction data on 1018 species pairs in four ecological groups (phytoplankton, bacteria/yeast, annual and perennial plants) to understand the drivers of coexistence. While the four ecological groups do not represent the diversity of natural systems, they do represent a variety of life spans, reproduction strategies, and habitats. We first quantified niche and fitness differences using one broadly applicable definition (J. W. Spaak and De Laender, 2020; J. W. Spaak, Godoy, et al., 2021). Next, we tested the hypothesis that niche differences (noted as 𝒩) *and* fitness differences (noted as ℱ) were larger *and* smaller, respectively, in coexisting species pairs, thus jointly promoting coexistence. We found that coexistence is mainly driven by niche and not fitness differences. We then carried out a clustering analysis in niche and fitness differences space to test for generalities across communities. This procedure identified two distinctly segregated clusters (each representing 40% of the data) that were only driven by niche differences and not by ecological group membership or methodological differences. We conclude that, for the four ecological groups considered, coexisting species differ from non-coexisting species because they have higher niche differences. Additionally, we conclude that there is broad similarity across the inspected communities, as they can be clustered in two groups only. The sharp boundary between and heterogeneity within these groups hints at the existence of unrecorded factors driving niche differences.

## 2 Methods

### 2.1 Data collection

We searched the literature for experimental measurements of niche and fitness differences (see Fig. S1 for an overview). To do so, we first identified eleven papers that have introduced a definition of niche and fitness differences (See Appendix, Table S5) and gathered all papers that cited one of these eleven original definitions by the 14*t h* of December 2020. For the highly cited definitions, more than 100 citations (Chesson, 2000b; Adler, HilleRisLambers, et al., 2007; Chesson and Kuang, 2008), we refined the search with the following keywords: (“niche differences” OR “niche overlap” OR “stabilizing mechanisms”) AND (“fitness differences” OR “fitness “ OR “equalizing mechanism”) AND (“Experiment” OR “Data” OR “Field”) AND (“Competition” OR “Coexistence”). Only articles that measured niche or fitness differences experimentally using one of the eleven definitions were considered. Out of the eleven definitions, seven were used empirically (Chesson, 2000b; Carroll et al., 2011; Godoy, Kraft, et al., 2014; Zhao et al., 2016; Saavedra et al., 2017; Bimler et al., 2018; J. W. Spaak and De Laender, 2020). We gathered a total of 639 papers, of which 50 contained experimental measurements of niche and fitness differences. These 50 papers contained 29 independent data-sets corresponding to 1018 communities (Appendix, section A.1, Table.S4). For each article, we extracted all species-specific growth parameters available (competition coefficient, sensitivity, intrinsic growth rate, invasion growth rate, etc.) and the outcome of species interaction, i.e. coexistence, competitive exclusion or priority effects. Additionally, we extracted ecological information about the community, such as the taxonomy of the competing species and the co-occurrence (sympatric or allosympatric), and methodological information about the experiment, such as the experimental setting (field, greenhouse or laboratory experiment), the community model fit to the empirical data (Lotka-Volterra model, Annual plant model or no model at all) and the method to measure species growth rates (field observations, growth rates over time or space for time replica, i.e. multiple plots with different initial abundances of competing species) (Appendix, section A.1, Table.S4). We labelled the taxonomy of the competing species into four different ecological groups, phytoplankton (170 communities), bacteria and yeast (128 communities) and terrestrial plants, which we subdivided into annual (459 communities) and perennial plants (261 communities). We grouped yeast with bacteria as we only found one study using yeast (Grainger, Letten, et al., 2019), this is justified by the size and habitat of both systems. Similarly, we split terrestrial plants into two subgroups because we had sufficient empirical data to do so.

### 2.2 Standardizing niche and fitness differences

For the 29 papers collected, the empirically measured niche and fitness differences were computed by different definitions and are therefore not directly comparable (Appendix, section A.2, Table.S5) (J. W. Spaak and De Laender, 2020; Godwin et al., 2020). To compare the different results we first converted them to the same model-independent definition of niche and fitness differences (J. W. Spaak and De Laender, 2020; J. W. Spaak, Godoy, et al., 2021). This definition was the only definition which was applicable to all datasets we have found. Many other definitions were not applicable, because they were designed for a specific community model, such as annual plant or Lotka-Volterra community models (Godoy, Kraft, et al., 2014; Chesson, 2018; Saavedra et al., 2017). Another widely used, model agnostic method, by Carroll et al., 2011 was not applicable because our data contained considerable amounts of facilitation.

To compute the definition of J. W. Spaak and De Laender, 2020 one needs the invasion growth rate *r*_*i*_, the intrinsic growth rate *µ*_*i*_, and the no-niche growth rate *η*_*i*_. The invasion growth rate is the growth rate of the focal species *i* when the resident species is at its carrying capacity 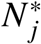. The no-niche growth rate is the growth rate of species *i* at the same converted density as species *j* ’s equilibrium density. This growth rate can be obtained via simulations. Given these three growth rates, they define niche differences as 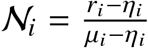 and fitness differences as 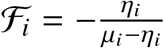. We use the updated notion for fitness as proposed by J. W. Spaak, Godoy, et al., 2021, as this leads to the coexistence condition of 𝒩_*i*_ > ℱ_*i*_. If only the invasion growth rate and the intrinsic growth rates are available, one can produce an estimate of the definition of J. W. Spaak and De Laender (2020) (Appendix, section A.2, Table.S5).

For 719 out of 1018 communities, the authors used a community model, such as the annual plant population dynamics (Levine and HilleRisLambers, 2009), which allows us to simulate all necessary growth rates and we therefore computed niche and fitness differences for these communities. For 234 out of 299 remaining communities, we found the invasion growth rates and the intrinsic growth rates, but not the no-niche growth rates. For these, we estimated niche and fitness differences (Appendix, section A.2). For the remaining 65 communities, we found the invasion growth rates and the carrying capacity, but not the no-niche growth rate nor the intrinsic growth rate. These were excluded from the analysis.

### 2.3 Clustering

We applied an automated clustering algorithm to the niche and fitness differences values of the inferior competitor of each community (species with ℱ_*i*_ > 0). We used an expectation-maximization algorithm with Gaussian kernels (mixture.GaussianMixture, EM clustering for Gaussian-mixture models) from the module ’sklearn’ version 0.23.2 in python version 3.8.5. The algorithm fits the best gaussian mixture to the data, i.e. each cluster consists of a location (mean of gaussian distribution) and a spread (covariance matrix of gaussian distribution). A species is not assigned to a specific cluster, but rather a probability is given that it belongs to each cluster (soft clustering). We first applied the clustering with one to ten clusters to identify the optimal number of clusters (Appendix A.3).

Next, we performed a meta-analysis using the package “metafor” (Viechtbauer, 2010) in R (R Core Team, 2020) to test whether the different clusters represented different ecological settings, e.g. ecological groups, experimental setting or used community model. With one exception, all communities from the same study had identical ecological and methodological information. With the `escalc()` function, we computed the proportion of species pairs belonging to each cluster within each study and across the empirical data. We use the sampling variances of those proportions as its precision. Then, with the `rma.uni()` function, we fitted linear mixed-effects models, with the estimated proportions of the clusters as effect sizes and with the different ecological settings used as qualitative moderators. If there are significant differences in the proportions of any of clusters for different ecological groups, experimental setting or community model, the moderator associated with this combination will be significantly different from zero, and the fitted mixed model will predict significantly different proportions for this combination.

## 3 Results

We identify three potential outcomes of species interactions: coexistence, exclusion, and priority effects. Overall, coexisting communities differed from other communities in their niche differences rather than in their fitness differences. We know that a persisting species *i* will satisfy 𝒩_*i*_ > ℱ_*i*_, while a non-persisting species *j* will satisfy 𝒩_*j*_ < ℱ_*j*_ (J. W. Spaak, Godoy, et al., 2021). By comparing these two species, we can therefore conclude that 𝒩_*i*_ ℱ_*i*_ > 𝒩_*j*_ − F_*j*_. However, from this inequality we can neither deduce 𝒩_*i*_ > 𝒩_*j*_ nor ℱ_*i*_ < ℱ_*i*_ ; we solely know that at least one of these must be correct. Depending on which of these inequalities is correct, 𝒩_*i*_ > 𝒩_*j*_ or ℱ_*i*_ < ℱ_*j*_, we may attribute coexistence to large niche differences or small fitness differences.

We found that in the empirical data, niche differences are mainly responsible for coexistence. In general, we have ℱ_*i*_ > ℱ_*j*_, but not necessarily ℱ_*i*_ < ℱ_*j*_. Across the 𝒩 - ℱ map, the communities were segregated along niche differences (Fig. 1 C) rather than the fitness differences (Fig. 1 A). Species from coexisting communities had significantly higher niche differences than species from other communities (median 0.864 and 0.019, Kruskal-Wallis *p* < 1*e* − 10). However, they differed not significantly in their fitness differences (median 0.362 and 0.411, *p* = 0.074). Species in communities with priority effects had lower niche and fitness differences than species from either coexisting or exclusion communities (medians -0.259 and 0.107, *p* < 1*e* − 5 in all comparisons). From this we conclude that, across the ecological groups considered, coexistence is driven by niche differences and not by fitness differences. Moreover, the values of niche and fitness differences capture both the high level-processes at play in the community (positive, negative or no frequency dependence, and facilitation) and the expected outcomes of these processes (competitive exclusion, coexistence, and priority effects). Here, negative density dependence was the most prevalent process (Fig. 1C), and mostly resulted in coexistence. Facilitation and positive frequency dependence mostly resulted in coexistence and exclusion, respectively.

**Figure 1:**
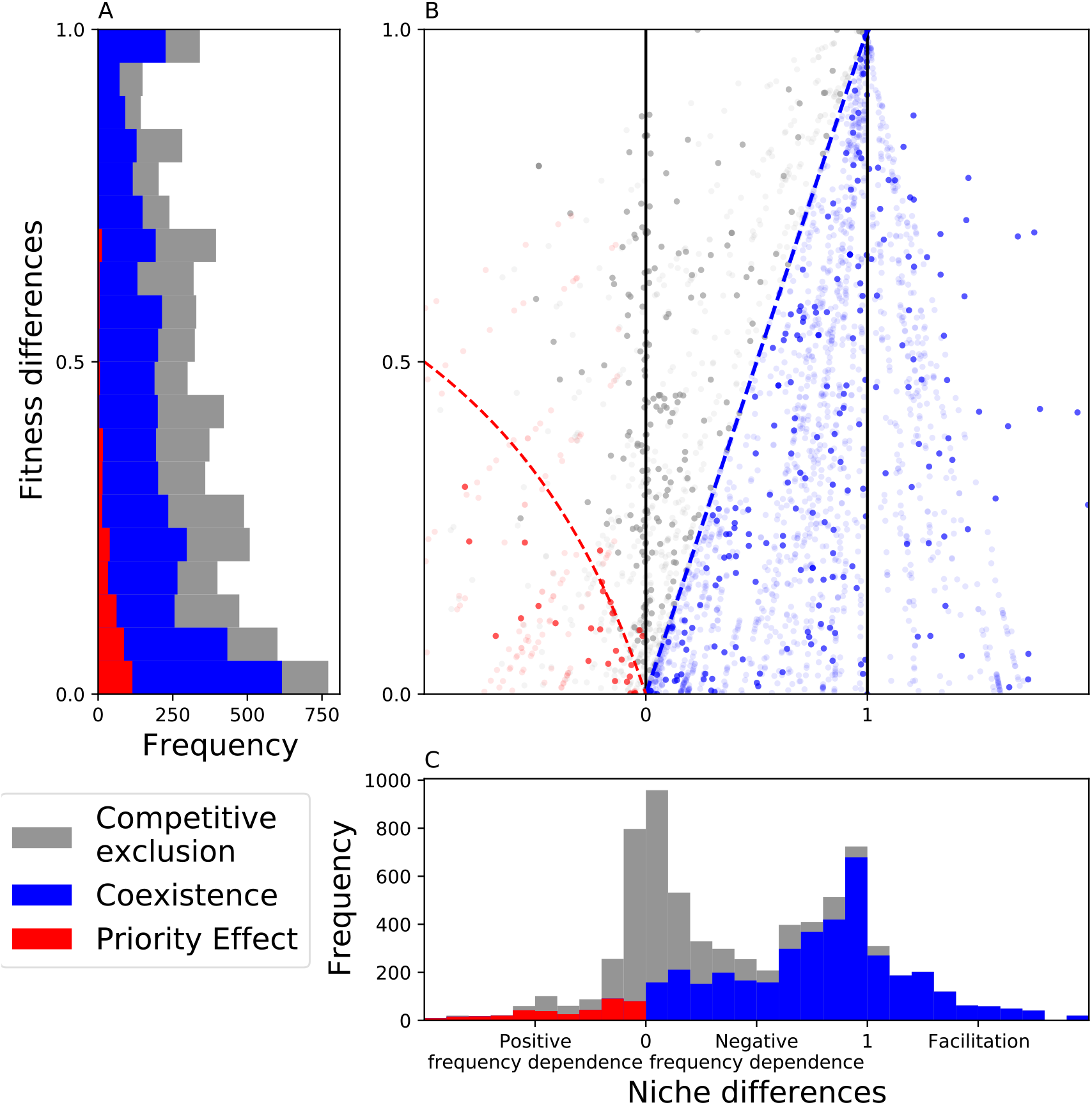
Niche and fitness differences of the inferior competitor for the analysed communities. A: The distribution of fitness differences for coexisting (blue) and non-coexisting communities (grey and red) are very comparable. Consequentially, fitness differences do not drive coexistence B: Distribution of all niche and fitness differences measured empirically. C: Species from coexisting communities (blue) have much higher niche differences than species from non-coexisting communities (grey and red). Additionally, species from communities driven by priority effects (red) have lower niche difference than species from communities driven by competitive exclusion (grey). We therefore conclude that coexistence is driven by niche differences. For some species we could only give an estimation of niche and fitness differences, for these, we plotted 10 random estimates (transparent points). The blue dashed line corresponds to the coexistence line, species below this line persist. The red dashed line delimits the region for priority effects if the community were driven by Lotka-Volterra dynamics. Species above this line can still belong to communities with priority effects, if their dynamics differ sufficiently from Lotka-Volterra dynamics.

**Figure 2:**
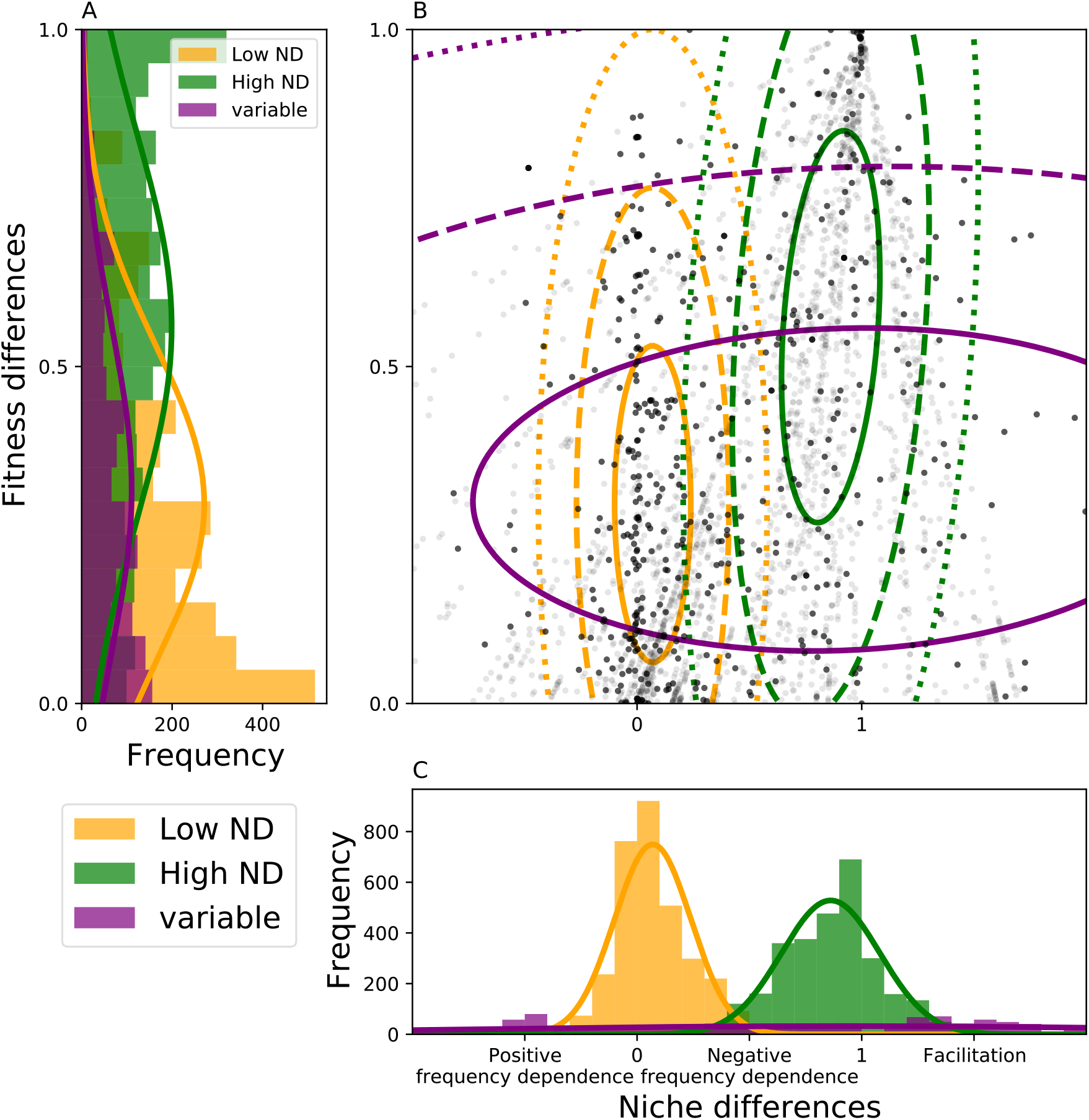
B: We clustered the niche and fitness differences into three clusters. The ellipses show one, two and three times the covariances, containing respectively 68, 95 and 99.7% of the data points within each cluster. The purple cluster contains less than 20% of the data (many outside the plotted range), the other clusters contain both about 40% of the data. A: Projection of the clusters to the fitness differences only, the fitness differences of the different clusters overlap substantially, indicating that fitness differences are not essential to the clustering. C: Projection of the clusters to the niche differences only. The green and orange cluster barely overlap, indicating that the sole knowledge of niche differences would be sufficient to cluster these two. We therefore conclude that clustering is driven by niche differences. The x-axis from panel A and y-axis from panel C differ from the corresponding panels in figure 1 because we do not stack the histograms in this graph, but did in the previous.

Nearly 80% of the data clustered into either of two distinct clusters (Fig. S3 B, Fig. 3 A, Appendix A.3). These two clusters correspond to the two previously observed peaks along the niche differences (Fig. 1C). We refer to them as “low 𝒩” (yellow) and “high 𝒩” (green). A third cluster, containing less than 20% of the data, represents a multitude of data points across the 𝒩-ℱ map and a much larger variance of niche differences. We therefore call this the “variable 𝒩” cluster (purple). Overall, the clusters had very similar fitness differences.

**Figure 3:**
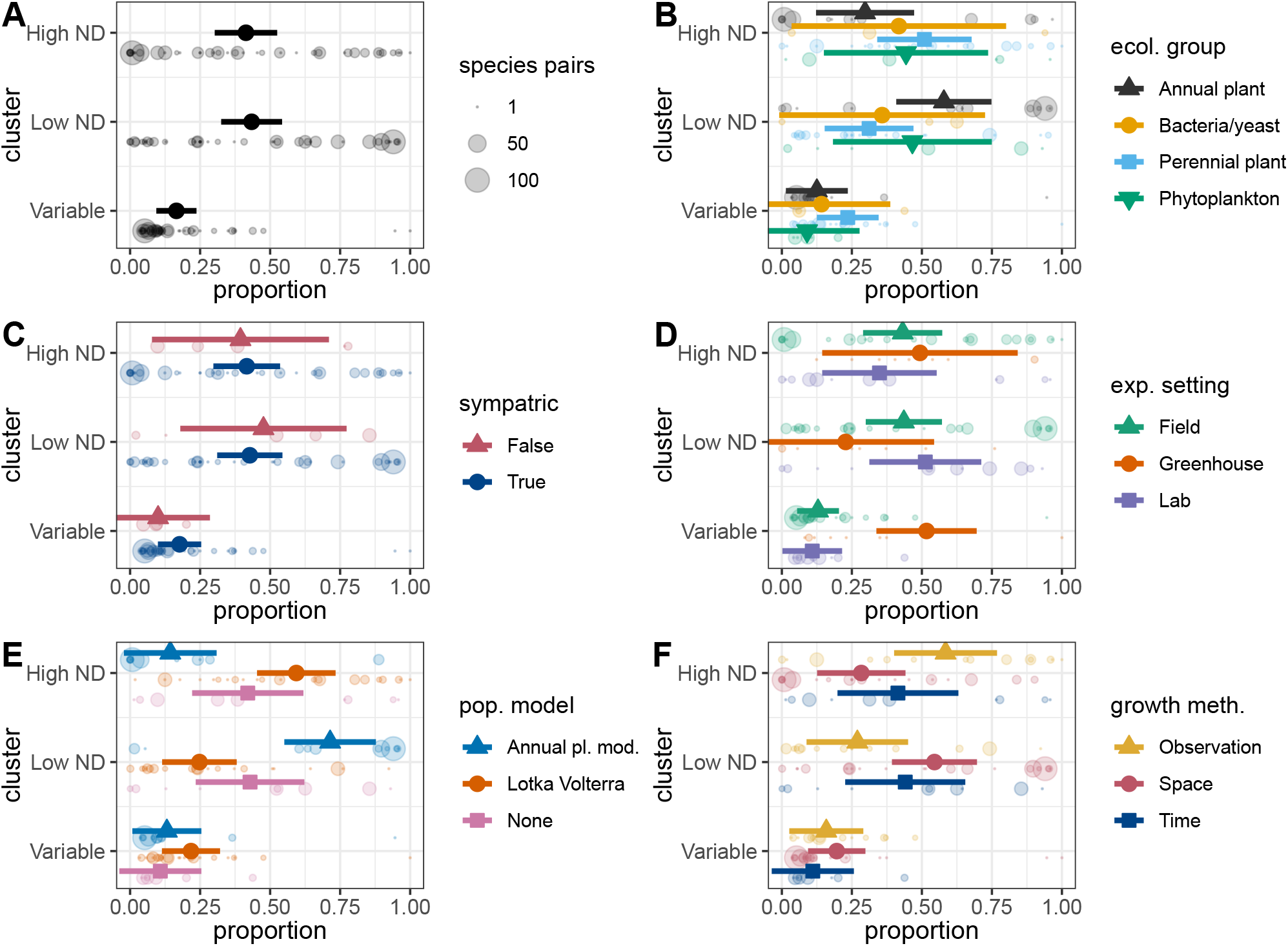
Proportion of species pairs belonging to the different clusters A–C (Fig. S3) obtained for different studies (semi-transparent points), and average proportions of the different clusters obtained through meta-analyses of the individual studies with error bars as confidence intervals (computed with the package “metafor” (Viechtbauer, 2010) in R (R Core Team, 2020)). In panel A, we represent the proportions obtained through random effects models of all the studies: 41 ± 6% of species pairs belong to cluster A, 44 ± 6% belong to cluster B, and 16 ± 4% belong to cluster C. B-F, proportions obtained through mixed-effects models considering respectively as moderators the ecological group of the species pairs (panel B), whether the species pairs are or are not sympatric (panel C), the experimental setting of the study (panel D), the employed population model (panel E), and the growth method (panel F, here divided between field observations, growth rates over time or space for time replica, i.e. multiple plots with different initial abundances of competing species). Generally these factors have no significant effect on the proportion of species pairs in the different clusters with the exception of greenhouse (panel D) and annual plant community model (Panel D). Thus, studies of different ecological groups, of sympatric or non-sympatric species, or with different experimental settings, population models, or growth methods would not differ in the proportion of species pairs that belong to each of the clusters.

The existence of mainly two distinct clusters suggests qualitative differences between both. For example, we might expect that a given cluster mainly contains species from a certain ecological group, e.g. annual and perennial plants, while the other cluster would be dominated by species from other ecological groups, e.g. phytoplankton and bacteria/yeast. However, this was not the case (Fig. 3 B). We similarly asked whether the co-occurrence (sympatric or asympatric), experimental setting (field, greenhouse or lab cultures), the used population model (Beverton-Holt annual plant model, Lotka-Volterra or no model at all) or how growth rates were assessed (using field observations, space for time replications i.e. multiple plots with different initial abundances of competing species, or growth rates over time) were driving these clusters. We found that, in general, the clusters did not differ significantly in any of these ecological or methodological aspects (Fig. 3 C-F). Rather, these clusters appear to be driven by inherent differences between coexisting and non-coexisting communities, which are consistent across these ecological and methodological differences.

## 4 Discussion

We used a model independent definition of niche and fitness differences for 1018 empirical species pairs, allowing for the first time a general analysis of species interactions across multiple ecological and methodological settings. We found that, first, coexisting communities have higher niche differences than non-coexisting communities (Fig. 1 C). Therefore, large niche differences, and not small fitness differences, drive species coexistence. Negative density frequency was also the most prevalent process. Second, we identified two main, clearly distinct, clusters, corresponding roughly to coexisting communities with high niche differences and non-coexisting communities with low niche differences (Fig. S3 B). Third, these clusters did not differ in ecological or methodological characteristics (e.g. ecological group, experimental setup, natural co-occurrence, population model used, and growth method). Instead, these clusters seemed to be consistent throughout all these differences.

### 4.1 Large niche differences, not small fitness differences, explained species coexistence at a local scale

Species coexist when niche differences exceed fitness differences. One could therefore expect that coexisting species have both higher niche differences and lower fitness differences than non-coexisting species. Yet, this hypothesis was rejected as we only found evidence for the former (Fig. 1). At a local scale, coexistence in the examined communities was primarily driven by mechanisms promoting niche differences, e.g. through self-limitation and net positive interactions (Hallett, Farrer, et al., 2018). These results consolidate and expand on findings from primary studies on annual plants (Levine and HilleRisLambers, 2009), perennial plants (Adler, Ellner, et al., 2010; Usinowicz et al., 2012), and phytoplankton (Narwani et al., 2013; Picoche and Barraquand, 2020).

The results also confirm earlier findings that niche differences are usually much stronger than necessary to coexist (Levine and HilleRisLambers, 2009; Chu and Adler, 2015), which is the case here for coexisting species from the high 𝒩 cluster. Our results show that niche differences are the main determinant for coexistence across ecological groups, highlighting the importance to sustain mechanisms that promote niche differences.

We also found that most coexisting communities were driven by negative frequency dependence (0 < 𝒩 < 1). Our results align with earlier findings in food web theory showing that negative frequency dependence should be present in up to 90% of interacting species (Barabás, Michalska-Smith, et al., 2017). In annual and perennial plants, intraspecific competition was previously found to be on average 1.5 and four to five times larger than interspecific competition, respectively (Adler, Smull, et al., 2018; Armitage and Jones, 2019). Yet, the prevalence of negative frequency dependence seems to expand beyond annual and perennial plants, across ecological groups, highlighting the importance of self-regulating mechanisms rather than species interspecific differences (Armitage and Jones, 2019). On the other hand, facilitation and positive frequency dependence were significantly less present. This complementary result supports that, not only mechanisms promoting niche difference, but specifically intraspecific mechanisms are important to maintain coexisting community.

The niche differences, and the corresponding processes we quantified (fig. 1.C) are the net result of multiple underlying mechanisms leading to multiple kinds of species interactions. Thus, the detection of specific interaction types (e.g. positive ones, Adler, Smull, et al. (2018) and Picoche and Barraquand (2020)) in a community does not guarantee specific processes (e.g. facilitation) will emerge in that community. A corollary of this is that our analysis gives no information about the prevalence of, for example, positive, asymmetric, or correlated species interactions – it only reflects the net result of such interactions. Thus, findings that positive interactions can be abundant in nature (Choler et al., 2001; Martorell and Freckleton, 2014; Soliveres et al., 2015; Wainwright et al., 2016; Bimler et al., 2018; Adler, Smull, et al., 2018; Picoche and Barraquand, 2020) do not contradict our result that net facilitation was only frequently observed in perennial plants and phytoplankton.

We have focused on species pairs, which by definition contain few species and a single interaction type per pair. We can therefore not be sure that fitness differences would continue to play a minor role for coexistence in communities with more species (Chu and Adler, 2015; Veresoglou et al., 2018; J. Spaak et al., 2021) inevitably hosting a more diverse set of species interaction types (Parmentier et al., 2020; Bartomeus et al., 2021), including asymmetric and higher-order interactions (Letten and Stouffer, 2019; Levine, Bascompte, et al., 2017; Mayfield and Stouffer, 2017). On the one hand, theory has shown that all else equal, species richness tends to not affect niche differences, while increasing fitness differences (J. Spaak et al., 2021). This would suggest that the latter will become more important for coexistence in more diverse communities. On the other hand, Godoy, Stouffer, et al. (2017) have proven the necessity of niche differentiation to permit coexistence in intransitive networks. Hence, a meta-analysis including multispecies communities is needed to explore the extension of our conclusions to more diverse communities.

### 4.2 Clustered niche differences

We found two main clusters in the niche and fitness differences map. These clusters can loosely be classified as low niche differences with non-coexisting communities (Fig. S3, orange) and high niche differences with coexisting communities (green). It is not surprising that the cluster with high niche differences will consist of mainly coexisting communities. However, the existence of such a clear clustering, the fact that it occurs predominantly along the niche differences axis, and its independence of ecological or methodological differences, are surprising. The heterogeneity within these clusters hints at generalities across ecological groups: communities with distinct ecology cannot be distinguished based on their niche and fitness difference signature. This finding is encouraging, as it implies that similar processes drive community dynamics in different species pairs. This information can help understanding biodiversity and predicting environmental impacts across a variety of systems (Soliveres et al., 2015). However, the result that ecological and methodological predictors could not explain these two clusters also reveals an important challenge: what drives clustering of high and low niche species pairs?

We offer two hypothesis to explain the existence of these clusters. The first is based on limiting similarity, i.e. that there is a limit to how similar coexisting species can be (Macarthur and Levins, 1967a; Meszéna et al., 2006). The coexistence of interacting species depends on the relative size of the niche space (the range of parameters where species have positive growth) and the niche width, the amount of niche space each species occupies. If the niche space is small relatively to the niche widths of the interacting species, then there is one optimal strategy which will competitively exclude all other species (Pastore et al., 2021; Kremer and Klausmeier, 2017; Barabás et al., 2016). In this scenario, the interacting species have small niche differences and would be part of the low 𝒩 cluster. Conversely, if the niche space is relatively large, many species will coexist with limited overlap, which would lead to high niche differences between interacting species, and consequently included them in the high 𝒩 cluster. In such a setting (large niche space), Pastore et al., 2021 performed a virtual evolution experiment of species along a one-dimensional resource axis (niche space) and found that coexistence is mainly driven by evolution towards large niche differences, which is also supported by previous work (Macarthur and Levins, 1967b; Slatkin, 1980; Stomp et al., 2004). Thus, interacting species should ultimately evolve towards one of these two extreme scenarios, creating two peaks on the niche axis. However, this hypothesis challenges recent findings, suggesting the predominant interaction of evolution with fitness differences rather than niche differences (Hart et al., 2019; Germain, Srivastava, et al., 2020; Pastore et al., 2021). The second hypothesis is based on lumpy coexistence, a combination between niche theory and neutral theory (Scheffer and Nes, 2006). Limiting similarity posits that species will eventually self-organize at equidistant positions along a one-dimensional niche axis (Macarthur and Levins, 1967a; Barabás et al., 2016; Meszéna et al., 2006). Lumpy coexistence describes the transient state before this optimal self-organisation. Species will exist in lumps of species with very similar traits and large gaps between these lumps of species. Species from different lumps will have dissimilar traits and large niche differences, corresponding to the high 𝒩 cluster. Conversely, species within a lump will have very similar niches and compete almost neutrally with each other, corresponding to the low 𝒩 cluster (Scheffer and Nes, 2006).

### 4.3 Limitations and perspectives

We present a cross-community synthesis of the processes driving coexistence. Like any meta-analysis, one can identify several limitations. First, our results are only valid for the ecological groups represented in the data, and other ecological groups may behave differently than the ones considered here. With few exceptions, the investigated communities consisted of basal species competing for abiotic resources. It is entirely possible that our findings do not apply to higher trophic levels, as research hints to the possibility that higher trophic levels are driven by different coexistence mechanisms (Shoemaker et al., 2019). A deeper understanding of how two-species coexistence links to multi-species coexistence might give us a better understanding of why niche differences are important in two-species communities. Second, we have little mechanistic understanding of why coexisting species exhibit higher niche differences leading to a prevalence of negative frequency dependence. Various mechanisms can be responsible for this result, including mechanisms driven by specific organismal traits (Gallego et al., 2019; Kraft et al., 2015), phenological traits (Godoy and Levine, 2014; Farrer et al., 2010; Martorell and Freckleton, 2014; Adler, HilleRisLambers, et al., 2009), and these mechanisms can be fluctuation-independent (Armitage and Jones, 2019) or fluctuation-dependant (Hallett, Shoemaker, et al., 2019). To provide such understanding here would be purely speculative, given our data.

Despite these limitations, the presented analysis suggests clustering of coexisting and non-coexisting species pairs, but at the same time a broad generality within these clusters. A logical next step is therefore to connect these results to biological insights into the considered community types (e.g. traits (Maire et al., 2012; Narwani et al., 2013; Kraft et al., 2015), or historical interactions (Germain, Weir, et al., 2016; Gilbert and Parker, 2016)). Doing so will contribute to a better understanding of the drivers of coexistence. ———–

## 5 Acknowledgement

We thank Dr. Oscar Godoy for comments on an earlier version of the manuscript. We thank all authors for kindly providing information and/or data from their studies.J.W.S received support from Schweizerischer Nationalfonds Early Post-doc mobility 470 (P2SKP3 194960). F.D.L. received support from grant of the Fund for Scientific Re471 search, FNRS (PDR T.0048.16).

