## Appendix for "Niche differences, not fitness differences, explain coexistence across ecological groups"

### A Appendix

#### A.1 Collection of data

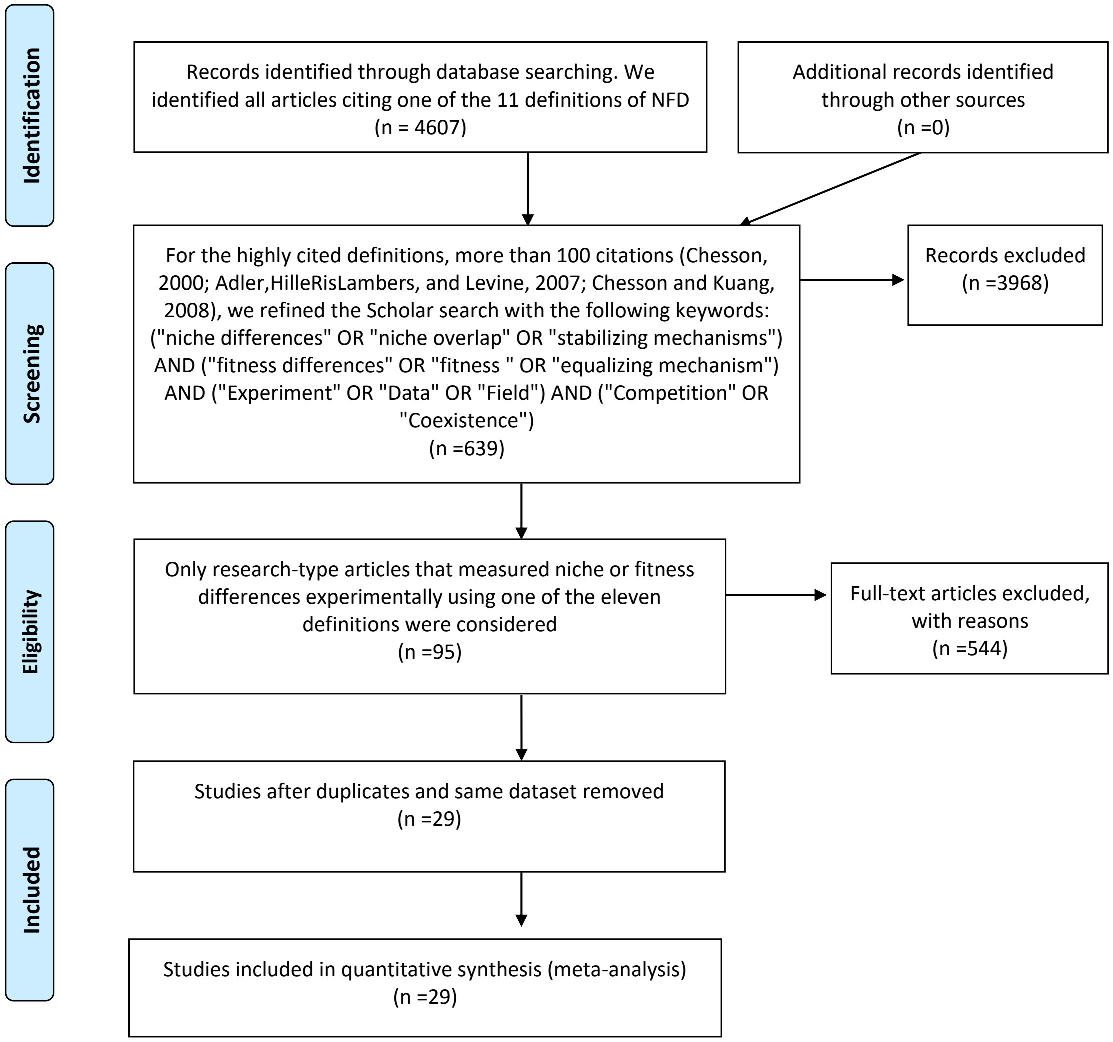

Figure S1: Flow-chart of meta-analysis.

| | Definition | Initial paper include data | Number of paper citing the definition until the $14^{th}$ of December 2020 | Refined if number of paper >100 | Empirical papers including the computation of $\mathcal{N}$ and $\mathcal{F}$ |
|---|---|---|---|---|---|
| | Godoy, Kraft, et al. (2014) | 1 | 96 | 96 | 23 |
| | Carroll et al. (2011) | 0 | 104 | 43 | 18 |
| | Adler, HilleRis-Lambers, et al. (2007) | 0 | 639 | 83 | 11 |
| | Chesson (1990b) | 0 | 3327 | 204 | 15 |
| | Spaak and De Laender (2020) | 1 | 2 | 2 | 0 |
| | Zhao et al. (2016) | 1 | 6 | 6 | 0 |
| | Bimler et al. (2018) | 1 | 20 | 20 | 6 |
| | Saavedra et al. (2017) | 0 | 58 | 58 | 5 |
| | Chesson (2003) | 0 | 83 | 83 | 6 |
| | Chesson and Kuang (2008) | 0 | 251 | 23 | 6 |
| | Carmel et al. (2017) | 0 | 10 | 10 | 1 |
| number total of papers: | 11 | 4 | 4596 | 628 | 91 |
| with the initial paper included: | | | 4607 | 639 | 95 |
| minus duplicate papers: | | | | | 50 |
| minus duplicate databases or non-available databases: | | | 41 | | 29 |

Table S1: The searching and refining processes lead us to use the data of 29 papers.

| | Field | Mesocosm | Greenhouse | lab | Total |
|---|---|---|---|---|---|
| Phytoplankton | 0.0 | 0.0 | 0.0 | 184.0 | 184.0 |
| Bacteria | 0 | 0 | 0 | 728 | 728.0 |
| Annual plant | 396 | 39 | 6 | 28 | 469.0 |
| Perennial plant | 215 | 0 | 18 | 28 | 261.0 |

Table S2: For each ecological group we report how many communities stem from which experimental setting. Phytoplankton and Bacteria have exclusively been analysed in laboratory settings. Similarly, annual and perennial plants have predominantly been analysed in a field setting.

| | Lotka Volterra | APM | AP2 | None | Total |
|---|---|---|---|---|---|
| Phytoplankton | 0.0 | 0.0 | 0.0 | 184.0 | 184.0 |
| Bacteria | 0 | 0 | 0 | 728 | 728.0 |
| Annual plant | 34 | 412 | 12 | 11 | 469.0 |
| Perennial plant | 253 | 8 | 0 | 0 | 261.0 |

Table S3: For each ecological group we report how many communities stem have been fitted with which population model. Phytoplankton and Bacteria have exclusively been analysed without a population model. Annual plants have predominantly been analysed with the Beverton-Holt annual plant model while perennial plants have predominantly been analysed with the Lotka-Volterra population model.

| Paper | Community type | Definition of $\mathcal{N}$ and $\mathcal{F}$ used | Population model used | Field or lab experiment | Number of 2 species communities | Sympatric | Growth method |
|---|---|---|---|---|---|---|---|
| Bimler et al. (2018) | Annual plant | Bimler et al. (2018) | AP2 | field | 12 | True | Obs |
| Godoy and Levine (2014) | Annual plant | Godoy, Kraft, et al. (2014) | APM | field | 153 | True | Space |
| Godoy, Kraft, et al. (2014) | Annual plant | Godoy, Kraft, et al. (2014) | APM | field | 15 | True | Space |
| Germain et al. (2016) | Annual plant | Godoy, Kraft, et al. (2014) | APM | field | 36 | NA | Space |

| | | | | | | | |
|---|---|---|---|---|---|---|---|
| Rey et al. (2017) | Annual plant | Godoy, Kraft, et al. (2014) | APM | field | 2 | True | Space |
| Hallett et al. (2018) | Annual plant | Godoy, Kraft, et al. (2014) | APM | field | 15 | True | Obs |
| Lanuza et al. (2018) | Annual plant | Godoy, Kraft, et al. (2014) | APM | field | 15 | True | Space |
| Matias et al. (2018) | Annual plant | Godoy, Kraft, et al. (2014) | APM | field | 45 | True | Space |
| Petry et al. (2018) | Annual plant | Godoy, Kraft, et al. (2014) | APM | field | 55 | True | Space |
| Wainwright et al. (2019) | Annual plant | Godoy, Kraft, et al. (2014) | APM | field | 24 | True | Space |

| | | | | | | | |
|---|---|---|---|---|---|---|---|
| Siefert et al. (2019) | Annual plant | Adler | APM | lab | 12 | | |
| Blackford et al. (2020) | Annual plant | Godoy, Kraft, et al. (2014) | APM | lab | 1 | True | Space |
| Armitage and Jones (2019) | Annual plant | Chesson (1990b) | Lotka Volterra | lab | 1 | True | Time |
| Ke and Letten (2018) | Annual plant | Chesson (1990b) | Lotka Volterra | lab | 3 | NA | NA |
| Ocampo-Ariza et al. (2018) | Annual plant | Carroll et al. (2011) | None | lab | 1 | False | Time |
| Zhang and Kleunen (2019) | Annual plant | Saavedra | APM | lab | 39 | True | Space |
| Tan et al. (2016) | Bacteria | Carroll et al. (2011) | None | lab | 7 | True | Time |

| | | | | | | | |
|---|---|---|---|---|---|---|---|
| Grainger et al. (2019) | Yeast | Carroll et al. (2011) | None | lab | 32 | True | Time |
| Li Shao-peng et al. (2019) | Bacteria | Carroll et al. (2011) | None | lab | 24 | True | Time |
| Zhao et al. (2016) | Bacteria | Zhao | None | lab | 65 | False | Time |
| Cardinaux et al. (2018) | Perennial plant | Godoy | APM | field | 8 | True | Space |
| Veresoglou et al. (2018) | Perennial plant | Chesson (1990b) | Lotka Volterra | lab | 28 | True | Obs |
| Adler, Smull, et al. (2018) | Perennial plant | Chesson (1990b) | Lotka Volterra | field and green-house | 255 | True | Obs and Space |
| Narwani et al. (2013) | Phytoplankton | Carroll et al. (2011) | None | lab | 27 | False | Time |

| | | | | | | | |
|---|---|---|---|---|---|---|---|
| Venail et al. (2014) | Phytoplankton | Carroll et al. (2011) | None | lab | 27 | False | Time |
| Gallego et al. (2019) | Phytoplankton | Carroll et al. (2011) | None | lab | 15 | False | Time |
| Jackrel et al. (2020) | Phytoplankton | Carroll et al. (2011) | None | lab | 99 | False | Time |
| Jia et al. (2020) | Phytoplankton | Carroll et al. (2011) | None | lab | 1 True | Time | |
| Spaak and De Laender (2020) | Phytoplankton | Spaak and De Laender (2020) | None | lab | 1 | True | Time |
| Total | 29 | | | | 1018 | | |

Table S4: Details of the 29 papers providing the data set used for the meta-analysis. Note that Adler, Smull, et al. (2018) is a meta-analysis gathering multiple papers. We have gathered ecological and methodological settings: the ecological group of the species pairs, the original definition of $\mathcal{N}$ and $\mathcal{F}$ used, the employed population model (Lotka Volterra, APM = Annual plant model; AP2 = Second annual plant model, that is the logarithm form of APM Bimler et al. (2018)), the number of pairwise community, whether the species pairs are or are not sympatric (true or false), the growth method divided between field observations (obs) , growth rates over time (time) or space for time replica (space), i.e. multiple plots with different initial abundances of competing species)

#### A.2 Computation of model independent method

We denote the monoculture growth rate as $\mu_i$, the invasion growth rate as $r_i$, the no-niche growth rate $\eta_i$ and the monoculture equilibrium density as $N_i^*$. With this we can define the three model independent definitions. For Spaak and De Laender, 2020 we define

$$\mathcal{N}_i = \frac{r_i - \eta_i}{\mu_i - \eta_i} \tag{S1}$$

$$\mathcal{F}_i = \frac{\eta_i}{\mu_i - \eta_i} \tag{S2}$$

The necessary parameters $\mu_i, r_i, \eta_i$ and $N_i^*$ where either reported in the original studies or the original papers reported enough information that allowed us to compute these parameters, most notably by reporting the per-capita growth rate function $f_i$. For time discrete per-capita growth rates functions (e.g. the annual plant model) we first log-transformed the growth rate, i.e. $\hat{f}_i(N_i, N_j) = \log(g_i(N_i, N_j))$.Given the per-capita growth rate function we can compute the monoculture growth rate as $\mu_i = f_i(0, 0)$, the monoculture equilibrium density as the solution of the equation $f_i(N_i^*, 0) = 0$ and the invasion growth rate as $r_i = f_i(0, N_j^*)$.

The only parameter that's left is the no-niche growth rate $\eta_i$, which is defined in

Spaak and De Laender, 2020. Numerically computing the no-niche growth rate $\eta_i$ is simple when using automated software (`https://github.com/juergspaak/NFD_definitions`). To do so, one has to solve the following two equations

$$c_i \cdot c_j = 1 \tag{S3}$$

$$\left|\frac{\mu_i - r_i}{\mu_i - f_i(c_j N_j^*, 0)}\right| = \left|\frac{\mu_j - r_j}{\mu_j - f_j(c_i N_i^*, 0)}\right| \tag{S4}$$

$\mu_i, \mu_j, r_i, r_j, N_i^*$ and $N_j^*$ can be computed before hand and can therefore be treated as constants in this equation. The only two variables are $c_i$ and $c_j$, whereof one can easily be eliminated with equation S3. Essentially, one therefore has to solve a one-dimensional equation. This equation might not be solvable by hand, but can easily be solved by numerical means. Importantly, if both species are strictly negative density dependence in monocultures, which is the case for most applications of modern coexistence theory, then the equation to solve is monotonic in $c_i$ or $c_j$ as well, which further simplifies numerical solutions.

Rather, the challenging part is to biologically justify the equations S3 and S4. $c_i$ converts density of species $i$ to densities of species $j$, and similarly $c_j$ converts densities of species $j$ to densities of species $i$. Consequently, $c_i \cdot c_j$ converts densities of species $i$ to densities of species $i$ and must therefore be one, which explains equation S3.

The $c$ are chosen such that they conserve total density, same total density implies same total effect on growth rates. $1 - \frac{r_i}{\mu_i}$ can be interpreted as the relative effect species $j$ has on the growth rate of species $i$. Similarly, $1 - \frac{\eta_j}{\mu_j}$ denotes the relative effect species $j$ has on itself. The total effect of species $j$ is therefore $\left(1 - \frac{r_i}{\mu_i}\right)\left(1 - \frac{\eta_j}{\mu_j}\right)$, which must be equal to the total effect of species $i$, which is $\left(1 - \frac{r_j}{\mu_j}\right)\left(1 - \frac{\eta_i}{\mu_i}\right)$. By reordering we obtain equation S4.

For the cases where the no-niche growth rate was not measured and could not be computed given the literature data we were able to approximate the definition of Spaak and De Laender, 2020 given the invasion and intrinsic growth rates $\mu_i$. First we note that for exactly one species $\eta_i$ is positive.

$$\eta_i = f_i(c_j N_j^*, 0) > 0 = f_i(N_i^*, 0) \tag{S5}$$

$$\Leftrightarrow c_j N_j^* < N_i^* \tag{S6}$$

$$\Leftrightarrow N_j^* < \frac{N_i^*}{c_j} = c_i N_i^* \tag{S7}$$

$$\Leftrightarrow \eta_j = f_j(c_i N_i^*, 0) < 0 = f_j(N_j^*, 0) \tag{S8}$$

Next we observe that

$$\eta_i < 0 \tag{S9}$$

$$\Leftrightarrow \mu_i - \eta_i > \mu_i \tag{S10}$$

$$\Leftrightarrow \frac{1}{\mu_i - \eta_i} < \frac{1}{\mu_i} \tag{S11}$$

$$\Leftrightarrow \frac{|\mu_i - r_i|}{\mu_i - \eta_i} < \frac{|\mu_i - r_i|}{\mu_i} \tag{S12}$$

We now assume, without loss of generality $\eta_i < 0$, which leads to

$$\frac{|\mu_i - r_i|}{\mu_i} > \frac{|\mu_i - r_i|}{\mu_i - \eta_i} = \frac{|\mu_j - r_j|}{\mu_j - \eta_j} > \frac{|\mu_j - r_j|}{\mu_j} \tag{S13}$$

Which is equivalent to

$$\frac{|\mu_i - r_i|}{\mu_i} > \frac{|\mu_i - r_i|}{\mu_i - \eta_i} > \frac{|\mu_j - r_j|}{\mu_j} \tag{S14}$$

$$\frac{\mu_i}{|\mu_i - r_i|} < \frac{\mu_i - \eta_i}{|\mu_i - r_i|} < \frac{\mu_j}{|\mu_j - r_j|} \tag{S15}$$

$$\mu_i < \mu_i - \eta_i < \frac{\mu_j}{|\mu_j - r_j|} \cdot (\mu_i - r_i) \tag{S16}$$

$$0 < -\eta_i < \frac{\mu_j}{|\mu_j - r_j|} \cdot |\mu_i - r_i| - \mu_i \tag{S17}$$

$$0 > \eta_i > \mu_i - \mu_j \frac{|\mu_i - r_i|}{|\mu_j - r_j|} \tag{S18}$$

$$\tag{S19}$$

We therefore chose 10 linearly spaced values for $\eta_i$ in the interval $[0, \mu_i - \mu_j \left|\frac{\mu_i - r_i}{\mu_j - r_j}\right|]$ and computed $\mathcal{N}_i$ and $\mathcal{F}_i$ based on these values.

Table S5: The 11 definitions, their frameworks and their restrictions. The symbols correspond to: $\alpha_{ij}$ interaction coefficients, according to Lotka-Volterra model; $a_{ij}$ interaction coefficients, according to the Annual plant model (Chesson, 1990a) ; $\lambda_i$ per-germinant fecundity in the absence of competition according to the Annual plant model ; $K_i$ environmental carrying capacity of species i; $k_{A1}$ is the per capita rate at which species A consumes resource 1.

| Paper | Assumption | mathematical definition | Experimental require-ments |
|---|---|---|---|
| Chesson and Kuang (2008) | Mac-Arthur model with predator extension | $\mathcal{N} = 1 - \frac{\sqrt{\alpha_{ij}^R \alpha_{ji}^R} + \sqrt{\alpha_{ij}^R \alpha_{ji}^R}}{s_i s_j}$ <br> $\mathcal{F}_i = \frac{s_j}{s_i} \frac{\mu_i^R - \mu_i^P - m_i}{\mu_j^R - \mu_j^P - m_j}$ | Model fitting |
| Chesson (1990b) | LV-model | $\mathcal{N} = 1 - \sqrt{\frac{a_{ij} a_{ji}}{a_{ii} a_{jj}}}$ <br> $\mathcal{F}_i = \sqrt{\frac{a_{ji} a_{jj}}{a_{ij} a_{ii}}}$ | Model fitting |

| | | | |
|---|---|---|---|
| Adler, HilleRis-Lambers, et al. (2007) | Annual plant model | $\mathcal{N}_i = \log\left(\frac{\lambda_j}{1+\frac{a_{ij}}{a_{jj}}(\lambda_j-1)}\right)$ $\mathcal{F}_i = \log(\frac{\lambda_i}{\lambda_j})$ | Model fitting |
| Godoy, Kraft, et al. (2014) | Annual plant model | $\mathcal{N} = 1 - \sqrt{\frac{a_{ij}a_{ji}}{a_{ii}a_{jj}}}$ $\mathcal{F}_i = \frac{\eta_j-1}{\eta_i-1}\sqrt{\frac{a_{ji}a_{jj}}{a_{ij}a_{ii}}}$ | Model fitting |
| Bimler et al. (2018) | Exponential Annual plant model | $\mathcal{N} = 1 - \sqrt{\frac{\exp a_{ij}+a_{ji}}{\exp a_{ii}+a_{jj}}}$ $\mathcal{F}_i = \sqrt{\frac{\exp a_{ji}+a_{jj}}{\exp a_{ij}+a_{ii}}}$ | Model fitting |
| Saavedra et al. (2017) | Linear species interactions | $\mathcal{N} = \frac{2}{\pi}\left(\arcsin\frac{\alpha_{ii}\alpha_{jj}-\alpha_{ij}\alpha_{ji}}{\sqrt{\alpha_{ii}2+\alpha_{ji}2}\sqrt{\alpha_{jj}2+\alpha_{ij}2}}\right)$ $\mathcal{F} = \frac{180}{\pi}\left(\arccos\frac{r.r_c}{\|r\|.\|r_c\|}\right)$ | Model fitting |
| Chesson (2003) | Invasion analysis possible, Population model available | $\mathcal{N} = \overline{\frac{\Delta I-\Delta N}{d}}$ $\mathcal{F}_i = \frac{r_i}{d_i} - \mathcal{N}$ | Model fitting |

| | | | |
|---|---|---|---|
| Carmel et al. (2017) | Invasion analysis possible, Minimal growth rate is equal to mortality | $\mathcal{N} = 1 - min(\frac{k_{A1}/k_{A2}}{k_{B1}/k_{B2}}, \frac{k_{B1}/k_{B2}}{k_{A1}/k_{A2}})$ $\mathcal{F}_i = min(\frac{k_{A1}k_{A2}}{k_{B1}k_{B2}}, \frac{k_{B1}k_{B2}}{k_{A1}k_{A2}})$ | Invasion growth rate, Mortality rate |
| Zhao et al. (2016) | Invasion analysis possible | $\mathcal{N} = 1 + r_i + r_j$ $\mathcal{F}_i = \log_{10}\left(\frac{K_i}{K_j}\right)$ | Invasion growth rate, Equilibrium abundances |
| Carroll et al. (2011) | Invasion analysis possible | $\mathcal{N} = 1 - \prod_{i=1}^{n} S_i^{1/n}$ $\mathcal{F} = exp\left[\left(\overline{(lnS)^2} - \overline{lnS^2}\right)^{1/2}\right]$ | Invasion growth rate, Monoculture growth rate |

| | | | |
|---|---|---|---|
| Spaak and De Laender (2020) | Invasion analysis possible | $\mathcal{N} = \frac{f_i(0,N_j^*)-f_i(c_jN_j^*,0)}{f_i(0,0)-f_i(c_jN_j^*,0)}$ $\mathcal{F}_i = \frac{f_i(c_jN_j^*,0)}{f_i(0,0)} - \frac{f_i(N_j^*,0)}{f_j(0,0)}$ | Density over time in mono-culture, Invasion growth rate |

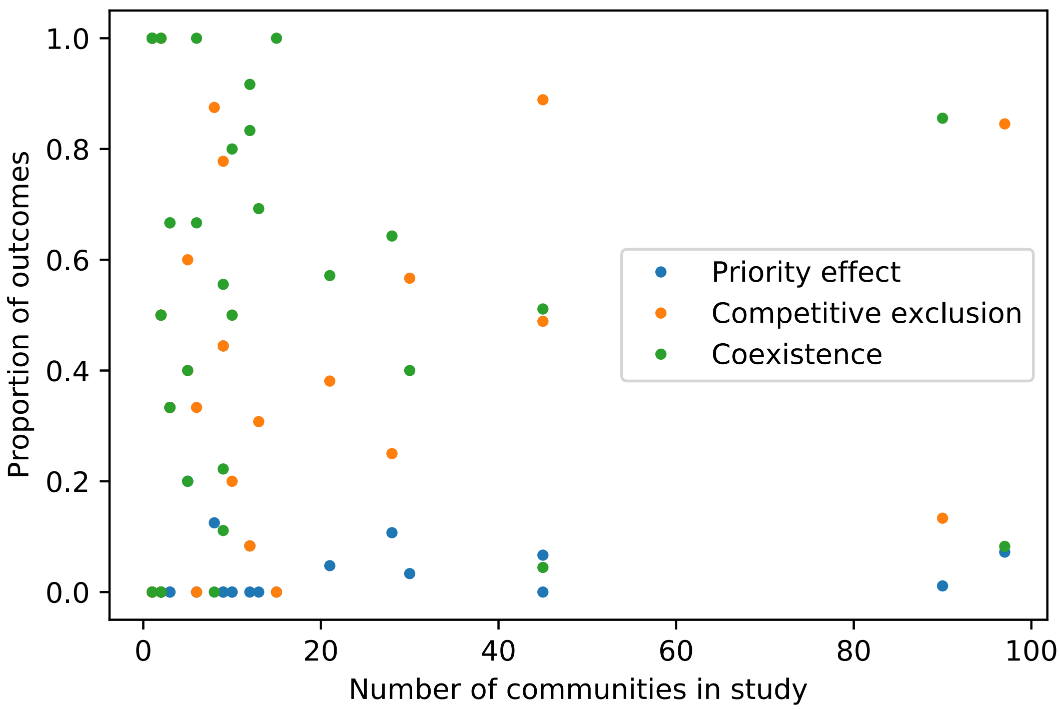

Figure S2: The size of the study (x-axis) did not have any effect on the probability of coexistence, priority effects or competitive exclusion. That is, the points are randomly spread along the both axis.

##### A.3 Clustering

We first analysed how many clusters there are in the dataset. To do so, we fit from one to ten clusters to the dataset, and replicated each fit 9 times. For each cluster we computed different metrics for goodness of fit. We used AIC, BIC, Log-likelyhood and the Rand-metric. The rand-metric compares for two given clusterings (with the same number of clusters) how often two data points have been clustered equally, i.e. whether they are in the same cluster in both clusterings or not. For AIC and BIC,

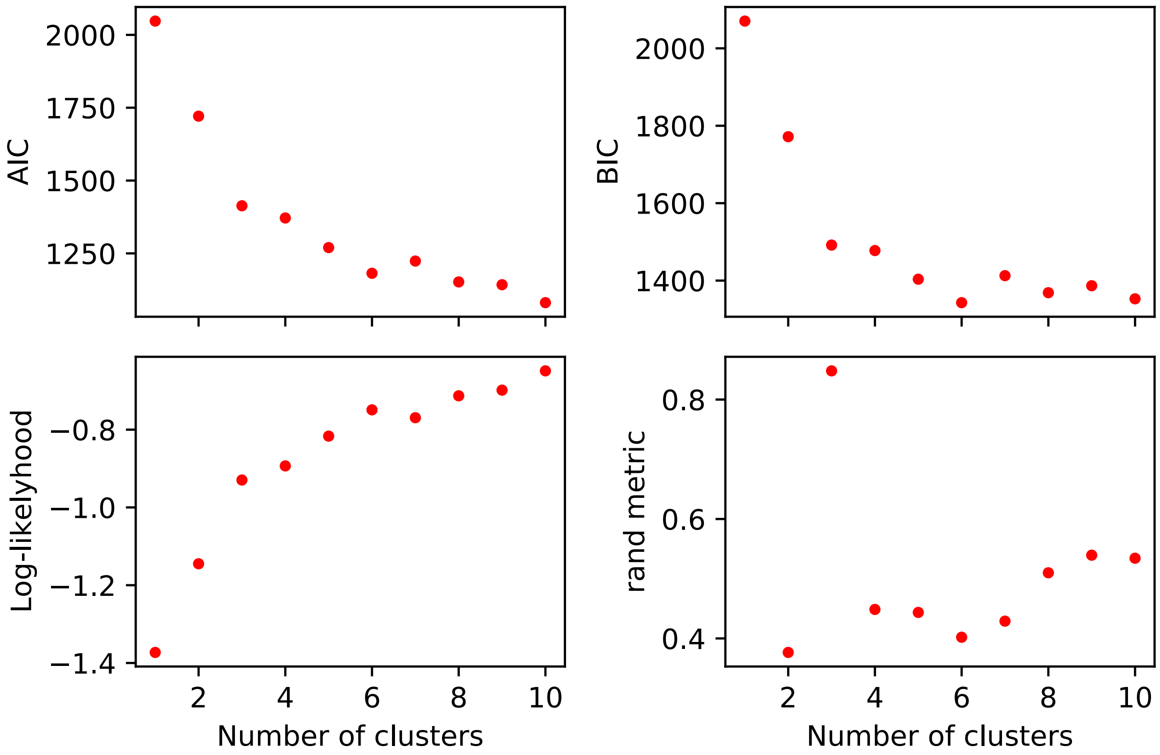

Figure S3: AIC, BIC, Log-likelyhood and rand metric all agree that 3 is the correct number of clusters for the data. AIC and BIC both initially decrease rapidly, unitl 3 clusters are reached. Adding more clusters only marginally decreases AIC and BIC. Similarly, Log-likelyhood initially increases much faster than after 3 clusters, again indicating that 3 is the correct number of clusters. Finally, Rand-metric is clearly highest for 3 clusters. The rand metric is not defined for only one cluster.

low values are better, for Rand metric and Log-likelyhood high values are better. Additionally, AIC, BIC and Log-likelyhood are known to increase with the number of clusters (similar to how model fit increases with increased number of variables). We therefore do not look at the absolute value of these, but at the relative change in these.
